# HIV protease inhibitor Saquinavir inhibits toll-like receptor 4 activation by targeting receptor dimerization

**DOI:** 10.1101/329003

**Authors:** Cheng Peng, Gang Deng, Yong Wang, Alzahrani Walid Abdulwahab, Hongwu Luo, Feizhou Huang, Hongbo Xu

## Abstract

Toll like receptor 4 (TLR4) is crucial in induction of innate immune response through recognition of invading pathogens or endogenous alarming molecules.Ligand-induced dimerization of TLR4 is required for the activation of downstream signaling pathways. TLR4 dimerization induces the activation of NF-kB and IRF3 through MyD88- or TRIF-dependent pathways. Saquinavir (SQV), a FDA-approved HIV protease inhibitor, has been shown to suppress the activation of NF-kB induced by HMGB1 by blocking TLR4-MyD88 association in proteasome-independent pathway. However, it remains nknown whether SQV is a HMGB1-specific and MyD88-dependent TLR4 signaling inhibitor and which precise signaling element of TLR4 is targeted by SQV. Our results showed that SQV inhibits both MyD88- and TRIF-dependent pathways in response to LPS, a critical sepsis inducer and TLR4 agonist, leading to downregulation of NF-kB and IRF3. SQV did not suppress MyD88-dependent pathway triggered by TLR1/2 agonist Pam3csk4. In the only TRIF-dependent pathway, SQV did not attenuate IRF3 activation induced by TLR3 agonist Poly(I:C). Furthermore, dimerization of TLR4 induced by LPS and HMGB1 was decreased by SQV. These results suggest that TLR4 receptor complex is the molecular target of SQV and shed light on that TLR4-mediated inmune responses and consequent risk for uncontrolled inflammation could be modulated by FDA-approved drug SQV.

## Introduction

Sepsis is a multifaceted host systemic inflammatory response syndrome (SIRS) caused by infection, and could be significantly amplified by endogenous factors[1]. Sepsis and subsequent organ dysfunction is one of the leading causes of death in intensive care units (ICUs)[2]. The lipopolysaccharide (LPS) from Gram-negative bacteria is a well-established early driver of sepsis. LPS could be detected by both cell surface and intracellular receptors, known as Toll like receptor 4 (TLR4) and caspase-4/5/11, respectively[3-5]. Caspases- and TLR4- dependent release of danger-associated molecular patterns (DAMPs) into the extracellular space during LPS-induced sepsis contributes to uncontrolled inflammation and host mortality[3, 6].

High-mobility group box 1 (HMGB1), a ubiquitous nuclear and cytosolic protein, is a strong DAMP mediator of excessive inflammation in sepsis[7, 8]. Extracellular HMGB1, released in late phase of sepsis, also signals through TLR4 in macrophages to induce cytokine release[9]. Thus, inhibitors of HMGB1-TLR4 signaling may provide a broad therapeutic time window for sepsis. In previous study, our group identified HIV protease inhibitor (PI) saquinavir (SQV) could block HMGB1 driven TLR4-MyD88 activation in macrophages and be protective in murine sepsis model[10].

SQV, as the first FDA-approved HIV protease inhibitor (PI), has been used for prolonged periods without severe side effects in HIV seropositive subjects since 1995. Unexpectedly, SQV and other HIV PIs possess pleiotropic antitumor and immunomodulatory effects, via suppressing activity of several mammalian targets in a concentration-dependent manner, including matrix metalloproteinase 2, 20S, 26S proteasome and akt phosphorylation[11]. However, it remains unknown whether SQV is a HMGB1-specific and MyD88-dependent TLR4 signaling inhibitor and which precise signaling element of TLR4 is targeted by SQV. Therefore, the present study investigated the molecular mechanism of SQV exerting anti-inflammatory property and modulating TLR4 activation.

## Results

### LPS-induced TLR4-MyD88 activation was inhibited by Saquinavir

Both LPS and disulfide HMGB1 signal through TLR4/MD2/CD14 complex at cell membrane level[12, 13]. In previous study, we reported SQV (Fig. 1A) blocks HMGB1-induced TNF-α production in human and mouse macrophages[10]. Here we tested whether SQV is a HMGB1-specific TLR4 inhibitor by evaluating the impact of SQV on LPS triggered TLR4 activation in cultured THP-1 monocyte-derived macrophages. SQV was found to potently suppress TNF-α release in THP-1 macrophages in response to LPS (Fig. 1B), while the compound control, another first-line HIV PI Atazanaivr didn’t affect LPS induced TNF-α production. Besides, in HEK293 cell transfected with TLR4/CD14/MD2 and NF-κB SEAP reporting system, NF-κB activation was significantly down-regulated by SQV (Fig. 1C). As shown before, HMGB1-TLR4 activated MyD88 pathway (IRAK4 phosphorylation and IRAK1 degradation) could also be prevented by SQV[10]. To determine whether SQV diminishes TLR4-MyD88 activation in a ligand-independent manner, we assessed degradation of IRAK1 in human macrophages exposed to LPS in the presence or absence of SQV. LPS stimulation led to the loss of IRAK1, but SQV attenuated LPS induced IRAK1 degradation and consequent IKK α/β phosphorylation (Fig. 1D). These results support the conclusion that SQV suppresses both LPS and HMGB1 induced TLR4-MyD88 activation.

**Figure 1.**
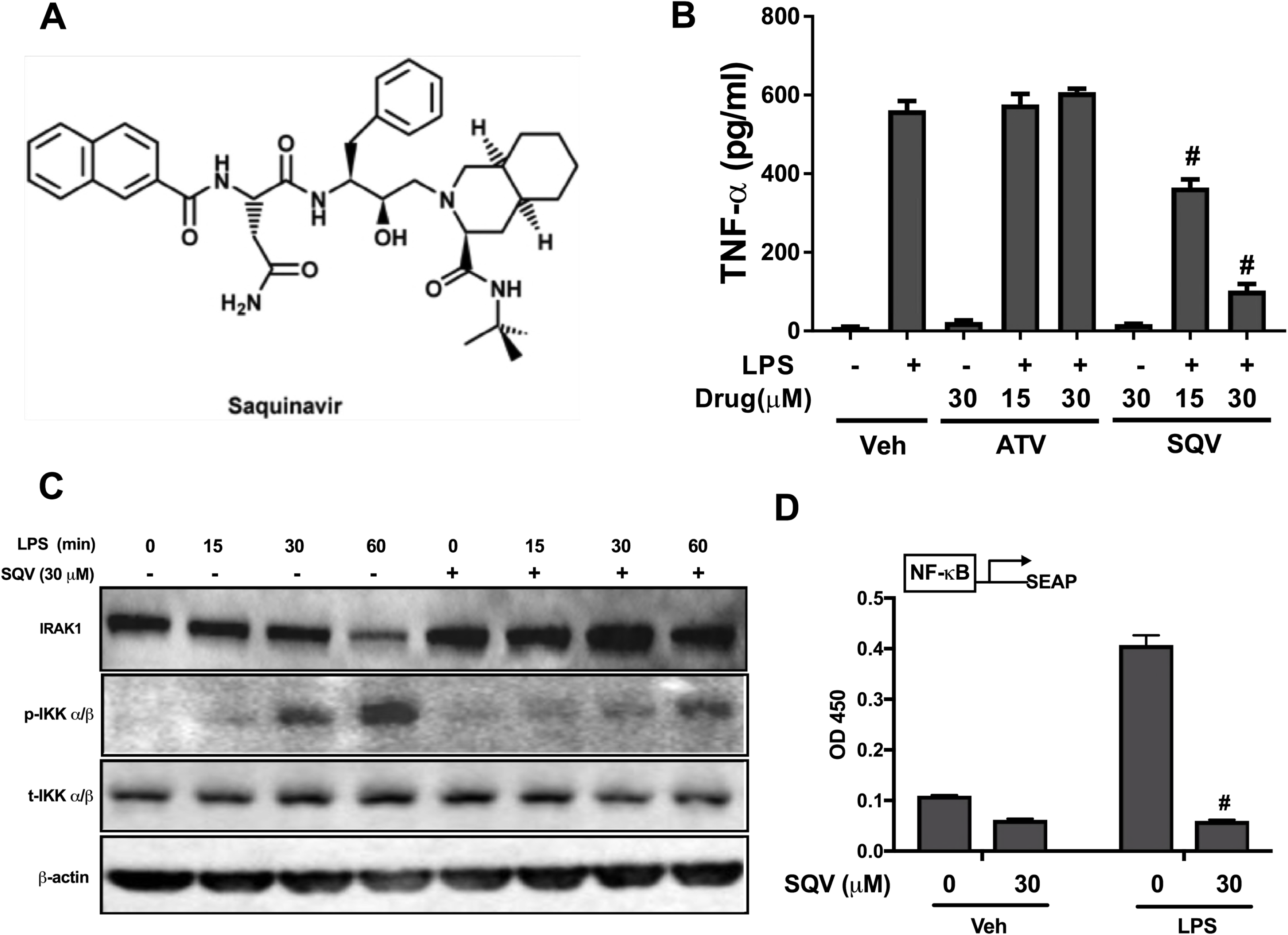
LPS-induced TLR4-MyD88 activation was inhibited by Saquinavir, but not Atazanavir. (A) The chemical structure of saquinavir. (B) PMA differentiated human THP-1 macrophages, pretreated with SQV or ATV (15, 30 µM) for 1 hour, were stimulated with LPS (1 ng/ml) for additional 16 hours. Cell supernatant was subjected to TNF-α ELISA assay. Data are presented as Mean±SEM (n=3). # Significantly different from LPS alone, p < 0.01. (C) THP-1 macrophages were incubated with SQV (30 µM) for 1h and then treated with LPS (5 ng/ml) for 15, 30 and 60 minutes. Cell lysates were analyzed for IRAK-1, phospho-IKK α/β, total-IKK α/β and β-actin protein by western blot. (D) HEK293 cells stably transfected with TLR4, MD2/CD14 and NF-kB-SEAP plasmids were pretreated with SQV (30 µM) for 1 h, then cells were stimulated with media control and LPS (5 ng/ml) for 24 hrs. SEAP levels in cell media were detected as the OD at 630 nm by spectrophotometer. Data are presented as Mean±SEM (n=6). # Significantly different from LPS alone, p < 0.01. The panels are representative data from more than three independent experiments. Veh, vehicle; SQV, Saquinavir; ATV, Atazanavir.

### LPS-activated TLR4-TRIF pathway was blocked by SQV

Except MyD88, TLR4 signal pathway relies on the other critical adaptor protein TRIF, which is recruited by TLR4 after LPS induced TLR4 endocytosis into endosome[12, 14]. The TRIF-dependent pathway culminates in the activation of both IRF3 and NF-κB. IRF3 is phosphorylated and transferred to nucleus promoting interferon-β transcription[14]. LPS led to the phosphorylation of IRF3 when stimulating THP-1 macrophages for 60 mins (Fig. 2A), and this phosphorylation was unexpectedly diminished by SQV, without affecting total IRF3. More interestingly, in the presence of SQV, LPS induced TLR4 endocytosis was decreased, as well as THP-1 macrophage polarization (Fig. 2B). Together, these findings indicate that SQV exerts anti-inflammatory effect through both TLR4-MyD88 and TLR4-TRIF pathways.

**Figure 2.**
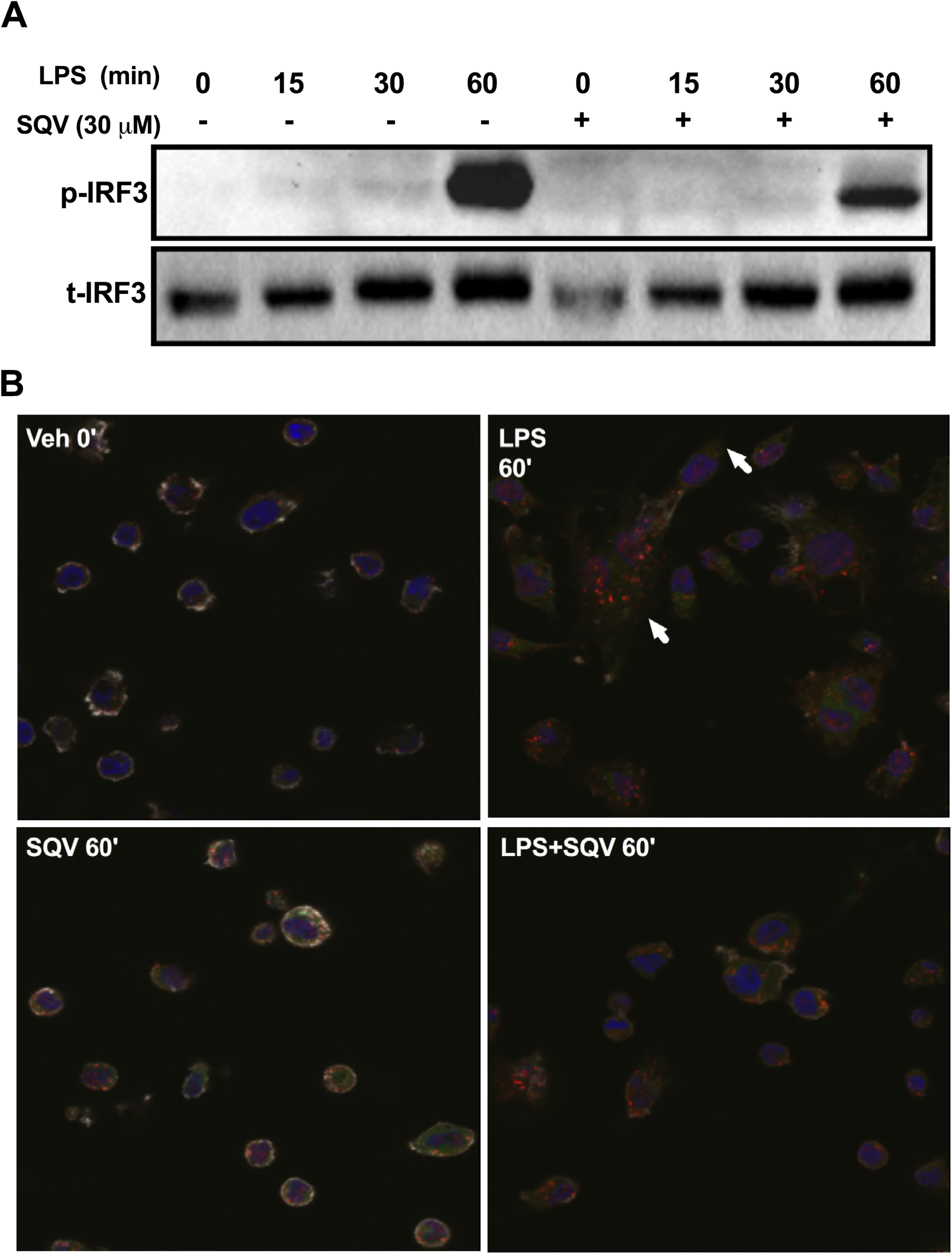
LPS-activated TLR4-TRIF pathway was blocked by SQV. (A) PMA differentiated human THP-1 macrophages were incubated with SQV (30 µM) for 1h and then treated with LPS (5 ng/ml) for 15, 30 and 60 minutes. Cell lysates were analyzed for phospho- and total-IRF3 protein by western blot. (B) PMA-differentiated THP-1 cells were stimulated with LPS (5 ng/ml) for 60 minutes after 1 h pretreatment of SQV (30 µM). Cell monolayers were fixed and stained for Cathepsin V (green) and EEA-1 (red). The panels are representative data from more than three independent experiments.

### SQV does not suppress the activation of MyD88 and TRIF induced by TLR 1/2 and 3 agonists

We couldn’t exclude the possibility that the targets of SQV lie in MyD88 or TRIF pathway. We were also curious about the selectivity of SQV on different Toll like receptors signaling. Therefore, we evaluated the impact of SQV on MyD88 and TRIF pathway when stimulated by other receptor-agonists engagement, using PMA-derived THP-1 macrophages. Since TLR1/2 is a MyD88-dependent cell surface TLR heterodimer[14], its specific agonist Pam3CSK4 induced phosphorylation of IRAK4 and degradation of IRAK1 (Fig. 3A and 3B), however these signal events were not affected even in the presence of SQV. TLR3 is the only TRIF-dependent and MyD88-independent TLR[14]. Poly (I:C), as the TLR3 agonist, triggered phosphorylation of IRF3, which was unaltered by SQV (Fig. 3C). Together, these results demonstrate that SQV specifically targets TLR4 receptor complex, rather than signal elements within MyD88 or TRIF pathways.

**Figure 3.**
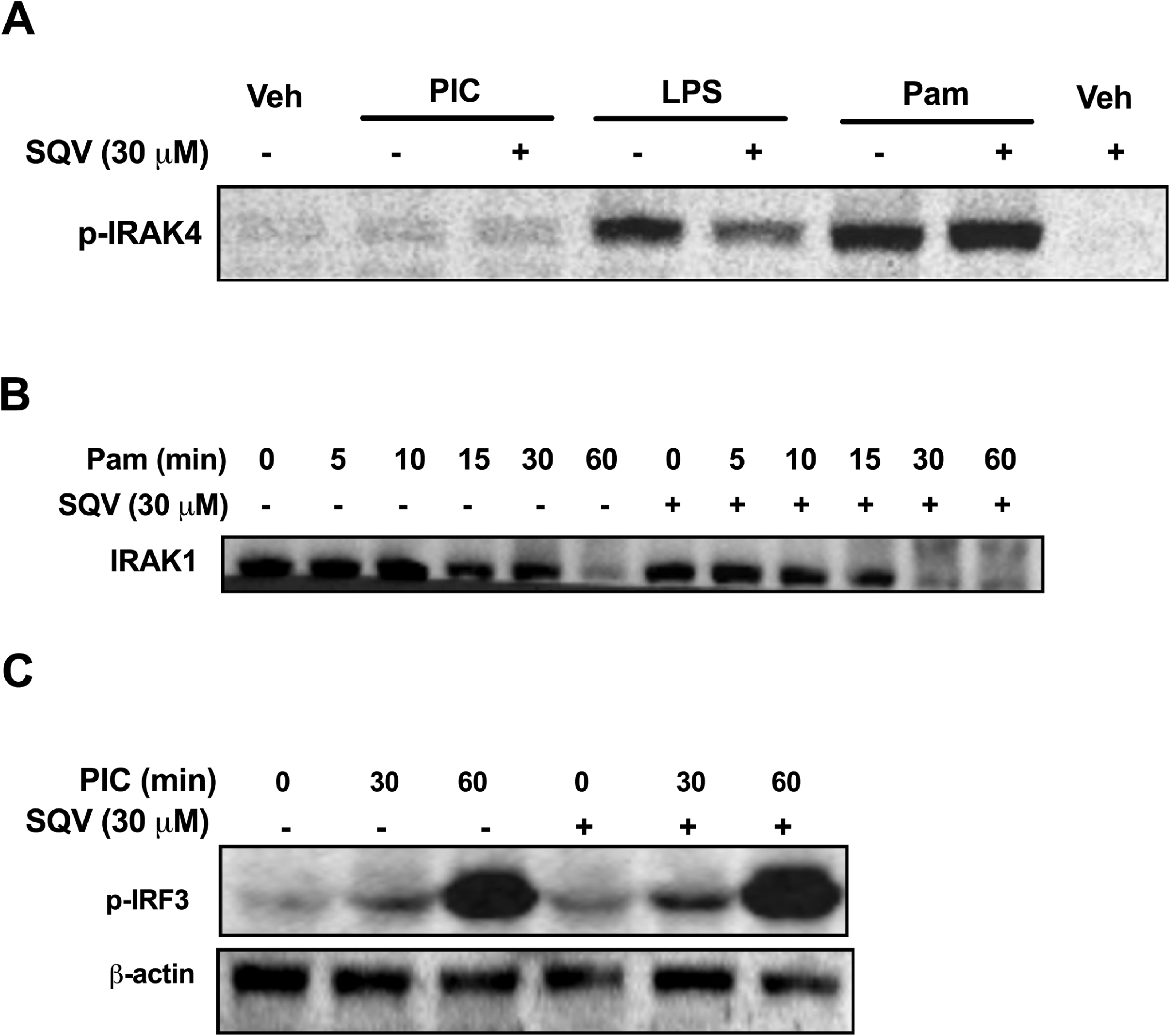
SQV does not suppress the activation of MyD88 and TRIF induced by TLR 1/2 andagonists. (A) PMA differentiated human THP-1 macrophages were pre-treated with SQV (30 µM) for 1h and then stimulated with Poly(I:C) (10 µg/ml), LPS (5 ng/ml) or Pam3CSK4 (100 ng/ml), for 30 minutes. Cell lysates were analyzed for phospho-IRAK4 protein by western blot. (B, C) PMA differentiated human THP-1 macrophages were treated with Pam3CSK4 (100 ng/ml) (B) or Poly(I:C) (10 µg/ml) (C) after 1h SQV incubation. Cells were lysed at indicated time points and IRAK1 degradation (B), IRF3 phosphorylation and β-actin protein (C) were assessed by western blot. The panels are representative data from more than three independent experiments.

### SQV suppresses LPS and HMGB1 induced TLR4 dimerization

The dimerization of TLR4 was shown to be a prerequisite for the ligand-induced TLR4 activation and acts upstream of MyD88 and TRIF pathway[15]. Since SQV inhibits both MyD88 and TRIF activation induced by LPS, therefore we next determined whether SQV disrupts LPS-induced dimerization of TLR4. For these studies, we used HEK293 cells stably transfected with MD2, CD14 and HA tagged human TLR4 as described previously[10]. We evaluated the dimerization of TLR4 by co-immunoprecipitation of TLR4-HA and transiently transfected TLR4-Myc. The dimerization, which was enhanced at 30 mins by LPS stimulation, was suppressed by SQV, while SQV didn’t modify the expression level of TLR4-Myc or the baseline level of TLR4 dimerization (Fig. 4A). Similar result was observed when using a different tagged TLR4 as TLR4-GFP in the case of HMGB1 stimulation (Fig. 4B). To explore the possible binding site of SQV on TLR4, an in silico molecular docking experiment was performed. The potential binding site was displayed in Fig 4C, showing the hydrophobic P3 and P1’ groups of SQV were embedded into the grooves of TLR4, overlapping the highlighted residues (Red) involved in the core dimerization interface of TLR4[6]. Therefore, the anti-inflammatory effect of SQV may initially be due to its ability to block TLR4 dimerization.

**Figure 4.**
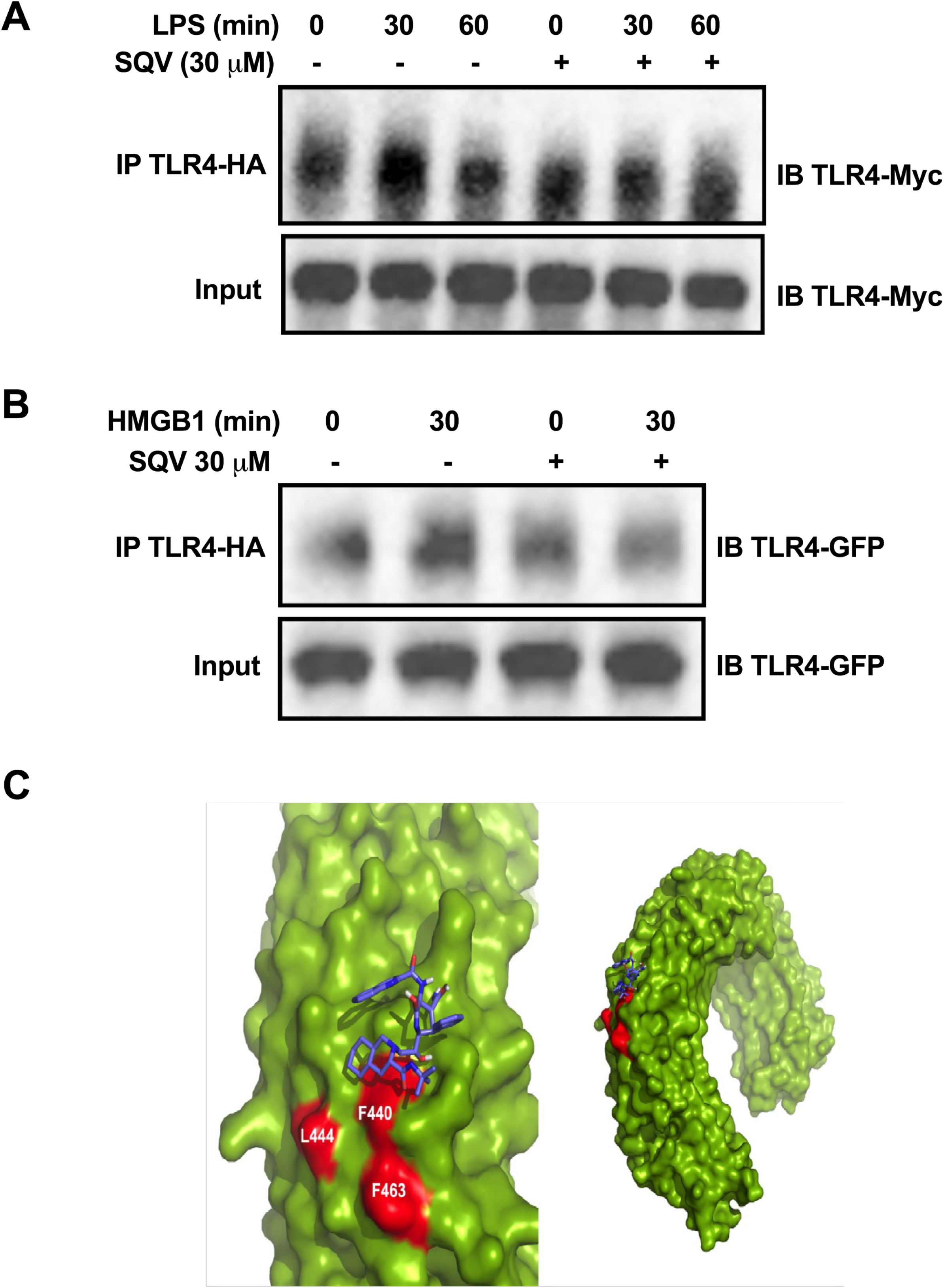
SQV suppresses LPS and HMGB1 induced TLR4 dimerization. (A, B) HEK293 cells expressing MD2/CD14, TLR4-HA and TLR4-Myc (A) or TLR4-GFP (B) were pretreated with SQV (30 µM) for 1h and then stimulated with LPS (5 ng/ml) or HMGB1 (5 µg/ml). Cells were lysed at indicated time points, then subjected to immunoprecipitation with anti-HA antibody and immunoblotted with anti-Myc (A) and anti-GFP (B) antibody. (C) The docking results of SQV with TLR4 complex: the whole and close-up view.

## Discussion

An increasing body of evidence highlights direct immunomodulatory effects of SQV in both experimental and clinical scenarios. Other than sepsis model, the preclinical data reported that SQV was protective in murine models of vascular intimal hyperplasia[16], liver ischemia-reperfusion (IR) injury[10], and lung Injury induced by liver warm IR[17] or mechanical ventilation[18]. In clinical aspect, a 6-month course of SQV was shown to reduce proteinuria and had a steroid-sparing effect in patients with nephrotic syndrome[19]. Recently, our colleagues proposed an exploratory clinical study to determine the safety and potential clinical benefit of the combination of HIV-PIs SQV and ritonavir in patients with idiopathic pulmonary arterial hypertension (IPAH)[20]. The basic mechanism of action is that SQV interact with human protein forming a drug-target complex, exerting off-label effects observed above. The mammalian target spectrum of SQV varies in a concentration-dependent manner. Additionally, based on current identified human targets of SQV, the novel targets may not definitely share similar structural properties as HIV proteases[11].

In this study, we provide the evidence that the anti-inflammatory mechanism of SQV arises from its direct modulation on TLR4-mediated signaling at the receptor level. TLR4 initiates immune response in the setting of infection and sterile inflammation through recognition of DAMPs and pathogen-associated molecular patterns (PAMPs). In sepsis scenario, LPS binding induces the formation of a symmetric M-shaped TLR4-MD-2-LPS multimer composed of two copies of the complex. Dimerization of TLR4 leads to the recruitment of MAL and MYD88 and the activation of the serine/threonine kinase IRAK4, which engages with the NF-κB and mitogen-activated protein kinase cascades, leading to the induction of pro-inflammatory cytokines[6]. TLR4 can also recruit TRIF (TIR domain-containing adaptor-inducing interferon-β) and TRAM (TRIF-related adaptor molecule) after endocytosis into endosomes, which leads to the activation of the transcription factor IRF3 and the induction of genes such as those encoding Type I interferons (IFNs)[14]. Therefore, suppression of TLR4 dimerization delete this scaffold for MyD88 or TRIF recruitment, thus preventing further signaling[21, 22]. Several studies have suggested that TLR4 dimerization could be a beneficial therapeutical target for anti-inflammation agents.

Our previous study happened to find, in human macrophages, SQV blocks HMGB1 activated TLR4-MyD88 pathway through inhibiting a cysteine protease cathepsin V other than proteasome. To current knowledge, TLR4 is the only toll like receptor signaling through both MyD88 and TRIF downstream pathway. LPS is the mostly investigated TLR4 ligand and a critical sepsis inducer. To evaluate therapeutical potential of SQV in sepsis, we tested the effect of SQV on LPS-triggered TLR4 signal pathway at both receptor and nuclear factor level. SQV, rather than control drug ATV, was found to inhibit LPS induced IRAK1 degradation and IKK α/β phosphorylation (Fig. 1C), as well as TNF-α (Fig. 1B) production and NF-kB (Fig. 1D) activation in human macrophages and HEK293 cell system. These findings confirmed that the target of SQV in TLR4 signaling lies upstream of IKK α/β and proteasome. Further probing on LPS-induced TRIF pathway showed that SQV blocks TLR4 endocytosis into endosome (Fig. 2B) and consequent IRF3 phosphorylation in response to LPS (Fig. 2A). These results suggest that SQV is a ligand-independent TLR4 inhibitor and could suppress both MyD88- and TRIF-dependent pathways in LPS-activated TLR4 signaling.

It is possible that SQV could target some components in MyD88 and TRIF pathways, respectively. Pam3CSK4 engages with TLR1 and TLR2 heterodimer, then the ligand-receptor recruit MyD88 from cell membrane and initates IRAK4 phosphorylation and IRAK1 degradation. Poly (I:C) is recognized by TLR3 homodimer at the endolysosome, followed by recruitment of TRIF and IRF3 phosphorylation. Pam3CSK4-TLR1/2-MyD88 and Poly (I:C)-TLR3-TRIF pathways share similar signaling components with LPS-TLR4-MyD88/TRIF pathways. However, both IRAK4 phosphorylation and IRKA1 degradation induced by Pam3CSK4 was not attenuated in the presence of SQV (Fig. 3A and B). Neither is Poly (I:C) triggered IRF3 phosphorylation (Fig. 3C). Together, these results indicate that the molecular targets for SQV include TLR4 receptor complex rather than downstream components in MyD88 or TRIF pathways.

Our immunoprecipitation results demonstrated that SQV could block both LPS and HMGB1 induced TLR4 dimerization (Fig. 4A and B), consistent with the hypothesis above. Futhurmore, docking analysis implies SQV may interact directly with F440 residue of TLR4, which is located on the TLR4 dimerization interface. The results of these experiments contribute to a novel mechanism for SQV modulating inflammatory response to TLR4 activation.

## Materials and methods

### Cell culture

Human monocytic THP-1 (ATCC) cells were differentiated for 12 h with 200 ng/mL PMA. HEK293 (ATCC), HEK293 stably transfected with CD14/MD2 and TLR4-HA or TLR4-Myc/GFP and human TLR4/NF-kB/SEAP reporter HEK293 cells (Invivogen) were grown at 37°C and 5% CO2, in DMEM (10% FBS, 2 mmol/L L-GLN, 100 U/mL PS, 1 mmol/L NaP, 100 µg/mL Normocin) (InvivoGen).

### Plasmids

pUNO-hTLR4-HA plasmid expressing the human TLR4 gene fused at the 3’ end to the influenza hemaglutinine (HA), pUNO1-hTLR4-GFP plasmid expressing the human TLR4 gene fused to a GFP gene and pDUO-MD2/CD14 plasmid coexpressing MD2 and CD14 genes (InvivoGen) were transfected into HEK293 cells using Genejammer transfection agent (Agilent) and Blasticidin (InvivoGen). pCMV6-hTLR4-Myc plasmid expressing the human TLR4 gene fused to a Myc gene was provided by Dr. Richard Shapiro.

### Reagents and Antibodies

OPTI-MEM (Invitrogen) was used as treatment buffer. Recombinant rat HMGB1, expressed in E. coli, was provided by Dr. KJ Tracey. LPS, Pam3csk4 and Poly(I:C) were from InvivoGen. PMA was a Sigma product. SQV and ATV were provided by Dr. Y Al-Abed. The following antibodies were used: anti-EEA-1 (NBP1-05962, Novus); anti-cathepsin V (MAB10801, R&D Systems); anti-HA (ab-hatag, InvivoGen); anti-IKK α/β(sc-7607, Santa Cruz); anti-phospho-IRF3(ab76493, Abcam); antibodies from Cell Signaling to IRAK1(4504), phospho-IRAK4 (11927), phospho-IKK α/β (2697), β-actin(4970), IRF3(11904), GFP(2956) and Myc-Tag(2278S).

### Immunoprecipitation and immunoblot

Cells were lysed in 1% NP-40, 50 mmol/L Tris-HCl (pH 7.4), 150 mmol/L NaCl, 10% glycerol and protease inhibitors (Sigma-Aldrich, Roche). After clearing by centrifugation, total protein content was quantified using a BCA kit and 1000 µg were immunoprecipitated (IP) overnight using anti-mouse/rabbit beads (Rockland) and corresponding antibodies. IP and input fractions were subjected to SDS-PAGE and Western blot analysis.

### NF-kappaB SEAP reporter assay

Human TLR4/NF-kB/SEAP reporter HEK293 cells stably transfected with MD2/CD14 plasmid were treated in HEK-Blue detection medium (Invivogen). SEAP levels in the medium were detected as the OD at 630 nm by spectrophotometer.

### Immunocytochemistry and immunoassay

PMA-differentiated THP-1 cells, seeded on coverlips, were fixed with 4% paraformaldehyde and then were made permeable by incubation in PBS containing 0.5% Triton X-100. After being blocked with 2% BSA, monolayer cells were stained for Cathepsin V (R&D MAB1080; green) and EEA-1 (Novus NBP1-05962; red). Fluorescence images were visualized with AxioVision 3.1 software (Carl Zeiss). Cell supernatants were assayed by enzyme-linked immunosorbent assay (ELISA) for TNF-α (R&D Systems).

### In silico molecular docking experiment

The TLR4-SQV docking studies were carried out based on the crystallography structures of human TLR4 structure (PDB: 3FXI). To prepare the receptor molecule for docking, all solvent molecules and the ligands in the structure were removed. The hydrogen atoms were added to the receptor structure. The grid maps was calculated by by AutoGrid v4.2 including the whole structure of a TLR4 monomer. The docking procedure was performed using AutoDock v4.2. The docking searches were not constrained in certain regions. The docked conformations of SQV were ranked by the lowest binding energy. Human TLR4 dimer interface was identified using PDBe PISA v1.52.

### Statistical Analysis

Data are expressed as means±SEM from at least three independent experiments. One-way analysis of variance (ANOVA) with Tukey post hoc test was used for comparison among all different groups. A P value of < 0.05 was considered statistically significant: #P < 0.01, unless otherwise indicated.

## ACKNOWLEDGMENTS

This study was supported by The New Xiangya Talent Project of the Third Xiangya Hosipital of Central South University (JY201519) and Provincial Natural Science Foundation of Hunan (2017JJ3419) to Hongbo Xu, and National Natural Science Foundation of China (81501791) to Yong Wang.

We are grateful to Drs. KJ Tracey, Y Al-Abed and Richard Shapiro for providing essential reagents and technical support.

